# Paladin is a PI(4,5)P_2_ phosphoinositide phosphatase that regulates endosomal signaling and angiogenesis

**DOI:** 10.1101/2020.02.11.943183

**Authors:** Anja Nitzsche, Riikka Pietilä, Chiara Testini, Takeshi Ninchoji, Ross O. Smith, Elisabet Ekvärn, Jimmy Larsson, Francis P. Roche, Isabel Egaña, Suvi Jauhiainen, Philipp Berger, Lena Claesson-Welsh, Mats Hellström

## Abstract

Cell signaling governs cellular behavior and is therefore subject to tight spatiotemporal regulation. Signaling output is regulated by specialized cell membranes and vesicles which contain unique combinations of lipids and proteins. The phospholipid phosphatidylinositol 4,5-bisphosphate, (PI(4,5)P_2_), an important component of the plasma membrane as well as other subcellular membranes, is involved in multiple processes, including signaling. However, which enzymes drive the formation and degradation of non-plasma membrane PI(4,5)P2, and their impact on cell signaling and function at the organismal level are unknown. Here we show in a mouse model that Paladin is a vascular PI(4,5)P_2_ phosphatase that regulates endosomal signaling and angiogenesis. Paladin was localized to the endosomal and Golgi compartments, and interacted with vascular endothelial growth factor receptor 2 (VEGFR2) *in vitro* and *in vivo*. Loss of Paladin resulted in increased internalization of the receptor, over-activation of extracellular regulated kinase, and hypersprouting of endothelial cells in the developing retina of mice. These findings suggest that inhibition of Paladin, or other endosomal PI(4,5)P_2_ phosphatases, could be exploited to modulate VEGFR2 signaling and angiogenesis, when direct and full inhibition of the receptor is not desirable.

## INTRODUCTION

In the eukaryotic cell, membranes in different subcellular compartments play distinct roles in cell signaling. Growth factor receptor signaling is initiated at the cell surface and continues after internalization and during endosome trafficking (Lampugnani, Orsenigo et al. 2006, Simons, Gordon et al. 2016). However, signaling is quantitatively and qualitatively distinct dependent on the specialized membrane compartment (Di Paolo and De Camilli 2006). Key to the maintenance of membrane specialization are lipid kinases and phosphatases that specifically phosphorylate/dephosphorylate phospholipids with an inositol head group i.e. phosphoinositides (PI). PIs are specifically distributed to generate ‘membrane codes’ on intracellular vesicles and the plasma membrane (Di Paolo and De Camilli 2006, Lemmon 2008). These PIs together with Rab GTPases are required for the maintenance and coordination of endocytosis and membrane trafficking (Jean and Kiger 2012) through recruitment of effector proteins to assemble specific endocytic complexes (Botelho, Efe et al. 2008, Jin, Chow et al. 2008, Lemmon 2008, Chagpar, Links et al. 2010, Mizuno-Yamasaki, Medkova et al. 2010). Consequently, as lipid kinases and phosphatases are key regulators of membrane identity and function, they are also regulators of cell signaling. However, the kinases and phosphatases involved in the generation of the specific PIs at distinct subcellular localizations are still not fully identified and their roles at the organismal level are only partially known.

PIs are phosphorylated at the 3′, 4′, and 5′ position of the inositol ring, giving rise to seven different PI species. The main PIs in the plasma membrane, early endosomes, late endosomes, and the Golgi apparatus are PI(4,5)P_2_, PI(3)P, PI(3,5)P_2_, and PI(4)P, respectively (Tan, Thapa et al. 2015). These PI pools present in microdomains of membrane vesicles provide a unique environment for signaling and sorting (Tan, Thapa et al. 2015).

Growth factor signaling is initiated at the plasma membrane, where the major PI pool is PI(4,5)P2 (Watt, Kular et al. 2002). PI(4,5)P_2_ is essential for signaling as a substrate for PI-3-kinase and the subsequent generation of the second messenger PI(3,4,5)P_3_ (Bilanges, Posor et al. 2019). PI(4,5)P_2_ is also present in intracellular membranes, as demonstrated by immune-EM and further suggested by the presence of lipid kinases and phosphatases, for which PI(4,5)P_2_ is as a substrate or product. An important role for PI(4,5)P_2_ dephosphorylation has been identified in growth factor receptor internalization and sorting in early endosomes. For example, PI(4,5)P_2_ generated by type I gamma phosphatidylinositol phosphate 5-kinase i5 (PIPKIγi5) regulates sorting of endosomal epidermal growth factor receptor (EGFR). PIPKIγi5-deficiency results in reduced transition of EGFR from endosomes to lysosomes and prolonged signaling (Sun, Hedman et al. 2013). However, a more general role for PI(4,5)P2 phosphatases has not been fully investigated.

Paladin is a membrane-associated protein encoded by *Pald1* or *x99384/mKIAA1274* in mouse and *PALD1* or *KIAA1274* in human. Its expression is primarily restricted to endothelial cells during development (Wallgard, Larsson et al. 2008, Suzuki, Moriya et al. 2010, Wallgard, Nitzsche et al. 2012). Although Paladin contains a phosphatase domain, it reportedly lacks protein phosphatase activity and was thus suggested to be a catalytically inactive pseudophosphatase (Huang, Hancock et al. 2009, Roffers-Agarwal, Hutt et al. 2012, Kharitidi, Manteghi et al. 2014, Reiterer, Eyers et al. 2014). However, Paladin has been implicated in various cell signaling pathways. A broad phenotypic screen in *Pald1 null* mice covering all organ systems revealed a specific lung phenotype, i.e. an emphysema-like lung histology and increased turnover of lung endothelial cells (Egana, Kaito et al. 2017). In addition, embryonic studies in chicken support a role for Paladin in neural crest migration (Roffers-Agarwal, Hutt et al. 2012). Cell culture studies suggest that Paladin negatively regulates expression and phosphorylation of the insulin receptor, as well as the phosphorylation of the downstream serine/threonine kinase AKT (Huang, Hancock et al. 2009). Furthermore, Paladin is a negative regulator of Toll-like receptor 9 (TLR9) signaling (Li, Wang et al. 2011). Collectively, these observations suggest that Paladin is an important player in cell signaling. Nevertheless, the mechanism whereby Paladin achieves those effects on diverse signaling pathways is unknown.

Here we provide evidence that Paladin is a novel PI(4,5)P_2_ phosphatase that lacks phospho-tyrosine/serine/threonine phosphatase activity. Paladin expression in endothelial cells regulates Vascular Endothelial Growth Factor Receptor 2 (VEGFR2) signaling at the level of endosomal trafficking. Vascular Endothelial Growth factor A (VEGF-A)/VEGFR2 signaling is essential for coordinating angiogenesis (Simons, Gordon et al. 2016). Consequently, genetic ablation of Paladin leads to retinal endothelial hypersprouting during development and excessive pathological angiogenesis. This suggests that targeting Paladin or other PI(4,5)P2 phosphatases could be explored to modulate VEGFR2 signaling.

## RESULTS

### Paladin is a PI(4,5)P_2_ phosphatase localized in endosomal vesicles

Paladin is considered to be a catalytically inactive pseudophosphatase. This is partially based on the sequence comparison to known protein phosphatases (PTP). The Paladin amino acid sequence contains four repeats of the minimal PTP consensus sequence CX_5_R (Figure 1a) (Wallgard, Nitzsche et al. 2012). Two of these repeats share high similarity with the extended conserved signature motif for PTP active sites, but importantly Paladin lacks the conserved histidine residue preceding the CX_5_R motif (Suppl Figure 1a) (Andersen, Mortensen et al. 2001). However, an increasing number of PTPs are recognized as being able to catalyze the dephosphorylation of phosphoinositides (Pulido, Stoker et al. 2013) and several new candidate phospholipid phosphatases have been proposed, including Paladin (Alonso and Pulido 2015). We therefore used a colorimetric screen based on the release of free phosphate from the substrate to evaluate such phosphatase activity of Paladin. We expressed and immuno-precipitated V5-tagged Paladin and phosphatase and tensin homolog (PTEN) proteins in HEK293 cells. Wild-type PTEN and dephosphorylation of PI(3,4,5)P_3_ was used as positive control and the C124S phosphatase-dead PTEN variant as negative control. Similarly, we used a Paladin variant with a Cys to Ser (C/S) substitution of all four cysteines in the CX_5_R motifs as a negative control (Suppl Figure 1a). Indeed, wild-type Paladin showed specific phosphatase activity towards PI(4,5)P_2_, and tended to also dephosphorylate PI(3,4,5)P_3_ but not PI monophosphates or inositol phosphates (Figure 1b and Suppl Figure 1b). Further, using radioactively labelled phosphopeptide substrate, with TC-PTP as a positive control, we confirmed the data by Huang and co-workers that Paladin lacks phospho-tyrosine activity (Figure 1c) (Huang, Hancock et al. 2009). Similarly, no Paladin activity against PKC-phosphorylated peptide was apparent (Suppl Figure 1c). These observations supported phospholipid phosphatase activity of Paladin.

**Figure 1.**
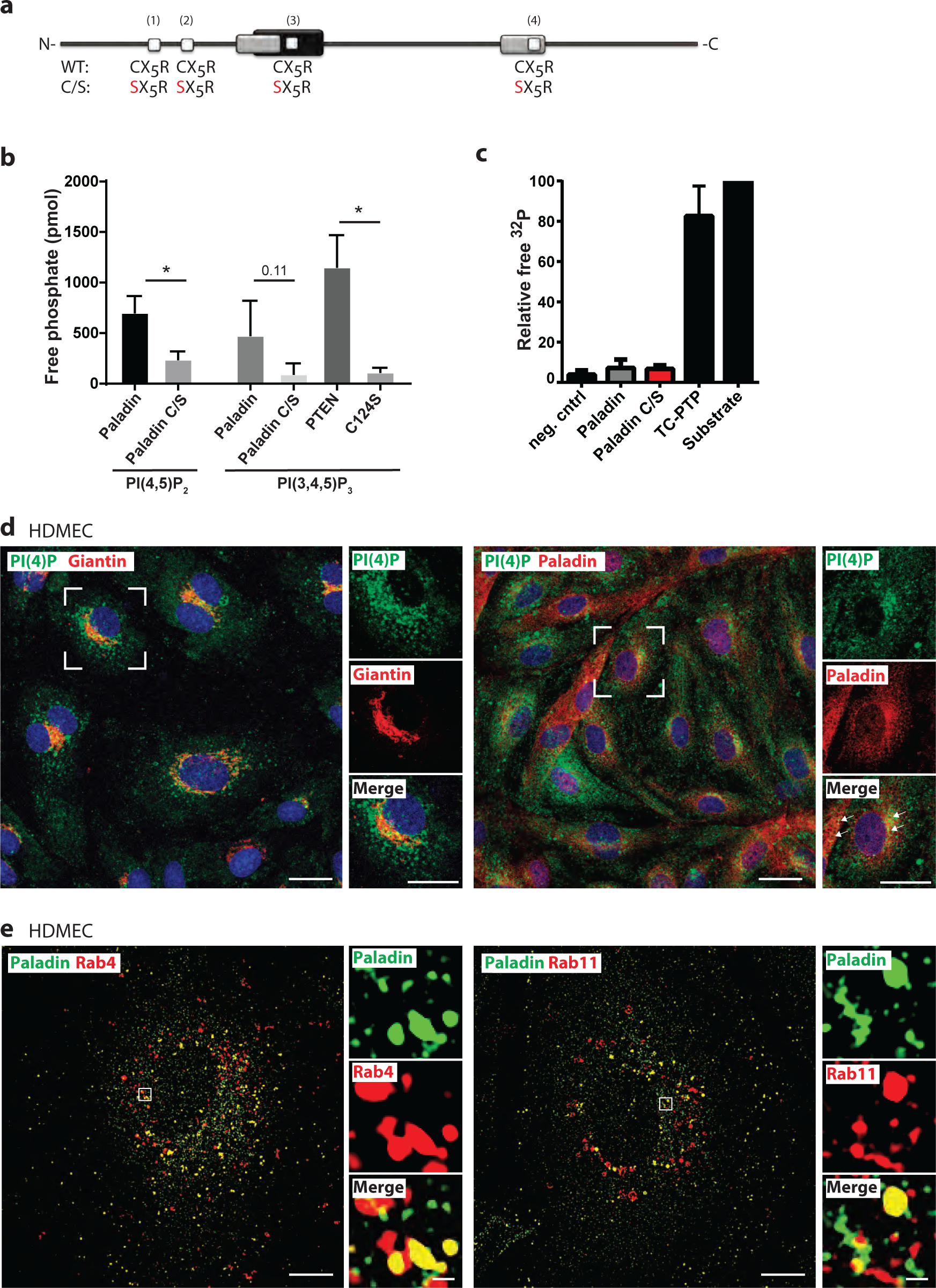
Paladin is a lipid phosphatase. **(a)** Schematic of Paladin protein depicting the four putative phosphatase domains (white boxes, CX_5_R; X: any amino acid; mouse: amino acids 121–127, 160–166, 315–321, and 664–670; human: amino acids 118–124, 157–163, 312–318, and 661–667). Phosphatase domains predicted by Interpro (black box) and SCOP SUPERFAMILY algorithm (grey). A full-length phosphatase-dead variant with the four cysteine residues substituted for serine (C/S). **(b)** Lipid phosphatase activity of Paladin, wild-type and phosphatase-dead C/S mutant, towards PI(4,5)P_2_ and PI(3,4,5)P_3_ substrates. Positive control; wild-type phosphatase and tensin homolog (PTEN); negative control; lipid phosphatase-dead C124S PTEN. Mean±SEM, unpaired *t*-test. n=3 individual experiments. **(c)** *In vitro* radioactive phosphatase assay using Paladin, wild-type or C/S variant, immunoprecipitated from HEK293, and as a substrate, Src-optimized peptide phosphorylated on tyrosines. Immunoprecipitates from cells transfected with empty vector or endogenous TC-PTP served as negative and positive controls. Data were normalized to ^32^P input. Mean±SEM. n=3. **(d)** Co-localization of Paladin and PI(4)P in HDMEC treated with VEGF-A–treated (50 ng/ml; 30 min) and immunostained for the presence of PI(4)P (green) and giantin (red), showing PI(4)P localization in the Golgi region (left panel). Analysis of non-treated HDMEC revealed partial co-localization of PI(4)P (green) and Paladin (red) (arrows in insert, right panel). Scale bars: 20μm. **(e)** Representative image of Paladin co-localization with Rab vesicles shown by super-resolution microscopy of VEGF-A–treated HDMEC (50 ng/ml; 10 min), immunostained for the presence of Paladin (green) and recycling vesicle markers Rab4 (red, left) or Rab11 (red, right), after transduction of cells with Rab4-cherry and Rab11-cherry baculovirus, respectively. Note the co-localization of Paladin and Rab4/Rab11 (yellow). Scale bars: 10 μm and 0.5 μm (inset). n=5 (Rab4) and 10 (Rab11).

Paladin is mainly expressed in endothelial cells during development (Wallgard, Nitzsche et al. 2012). Accordingly, we used immunostaining to evaluate the cellular location of Paladin in primary human dermal microvascular endothelial cells (HDMEC). The analysis revealed a vesicular staining pattern of Paladin but no staining of the plasma membrane and no co-localization with the junctional protein VE-cadherin was observed (Figure 1d,e and Suppl Figure 1d). We hypothesized that vesicular membrane localization could be facilitated by the N-terminal Paladin myristoylation, (Suzuki, Moriya et al. 2010), and that selective membrane localization could be enhanced by binding to phosphoinositides. We therefore tested the binding of Paladin to various phosphoinositides. We found that recombinant Paladin did indeed bind specifically to monophosphoinositides PI(3)P, PI(4)P, and PI(5)P, and PI(3,5)P_2_ (Suppl Figure 1e). This might reflect Paladin’s localization: e.g. PI(4)P is present in the Golgi membrane (Choudhury, Hyvola et al. 2005) and, accordingly, Paladin co-localized with PI(4)P and the Golgi marker giantin (Figure 1d and Suppl Figure 1f). This indicated that Paladin was localized to specific vesicular structures in the endothelial cells but not the plasma membrane.

To further characterize the subcellular localization of Paladin, we analyzed its localization in HDMEC expressing fluorescently tagged Rab4, −7, and −11, markers of fast recycling, slow recycling and late endosomes, respectively. Paladin co-localized mainly with Rab4 and Rab11. Super-resolution microscopy revealed that one-quarter of the Rab4- or Rab11-positive structures were also positive for Paladin, with Paladin seemingly localizing in sub-domains on Rab-positive vesicles (Figure 1e). Paladin also co-localized with Rab7-positive late endosomes, but not with EEA1 early endosomes (Suppl Figure 1g,h).

Taken together, Paladin catalyzes PI(4,5)P_2_ dephosphorylation but lacks phosphatase activity towards phosphorylated amino acid residues. Further, it co-localizes with the endosomal markers Rab4, −7, and −11, and Golgi membrane, but not the plasma membrane.

### Paladin regulates VEGFR2 internalization

Paladin is highly expressed in the vasculature, and regulates endothelial proliferation and survival in the lung (Wallgard, Nitzsche et al. 2012, Egana, Kaito et al. 2017). VEGF-A and its main receptor VEGFR2 are essential regulators of endothelial cell function and therefore we hypothesized that Paladin might interact with VEGFR2. We thus tested the effect of endothelial cell treatment with VEGF-A on Paladin and VEGFR2 localization *in vitro* and *in vivo*. We observed the formation of a Paladin-VEGFR2 complex in response to VEGF-A treatment both *in vitro* (in primary endothelial cells) and *in vivo* (in a mouse model). This interaction was further stabilized with blocking dephosphorylation by peroxyvanadate treatment (Figure 2a-c). Accordingly, super-resolution microscopy analysis confirmed VEGF-A induced co-localization of Paladin and VEGFR2 (Figure 2d). This indicated that Paladin could be involved in VEGF-A/VEGFR2 signaling.

**Figure 2.**
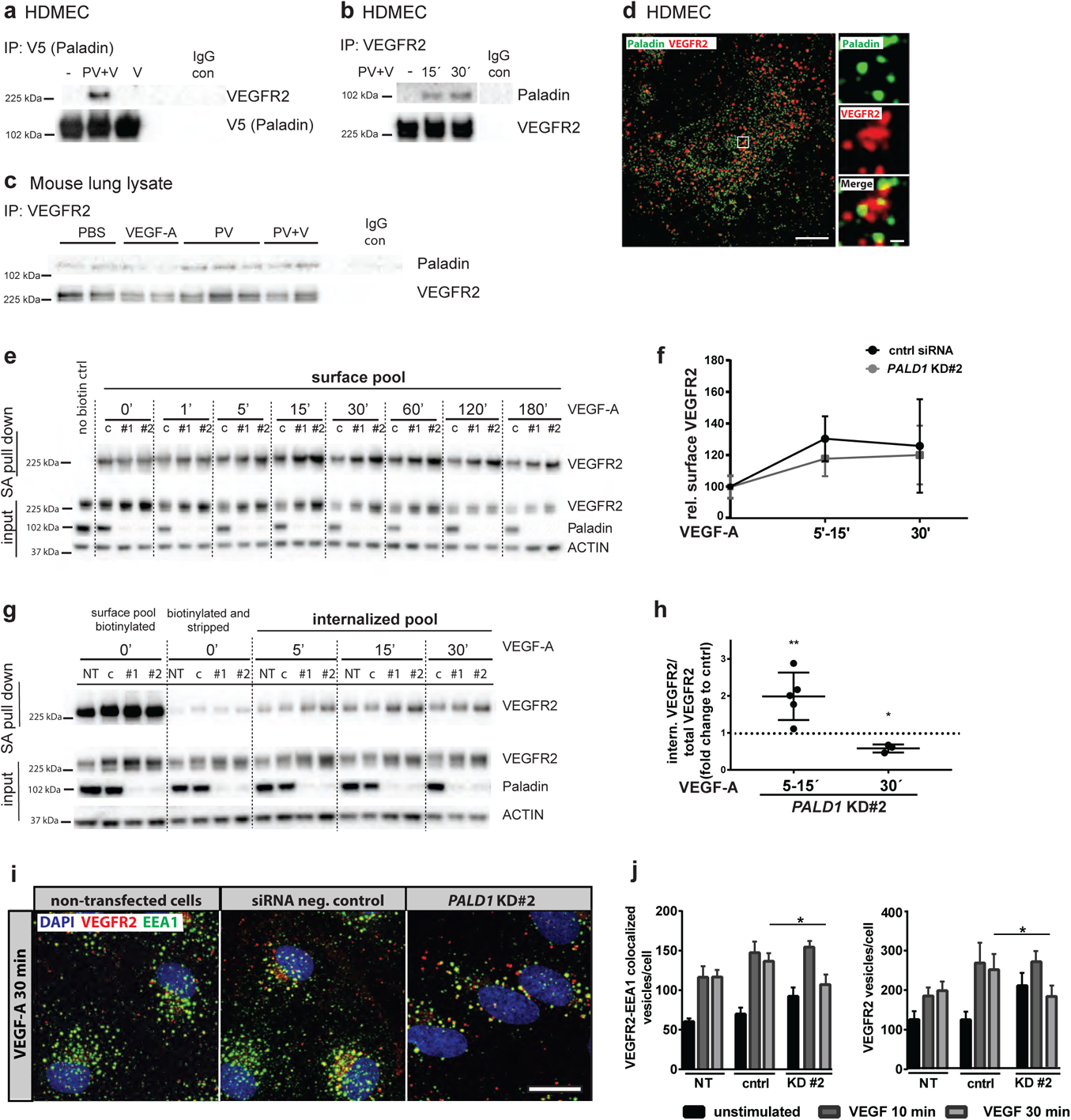
Paladin interacts with VEGFR2 and regulates its endosomal trafficking. **(a,b)** Formation of VEGFR2/Paladin complex *in vitro*. Immunoprecipitation (IP) of V5-tagged Paladin (a) or VEGFR2 (b) from *PALD1*-overexpressing (a) or untransfected (b) HDMEC stimulated with 50 ng/ml VEGF-A (V) alone for 30 min (a) or in combination with phosphatase inhibitor 100 µM peroxivanadate (PV) for 15 min (a,b,) and 30 min (b). Immunoblotting for Paladin, V5, and VEGFR2. IP with isotype control IgG as negative control (IgG con). **(c)** Formation of VEGFR2/Paladin complex *in vivo.* IP of VEGFR2 from lysate of adult mouse lung retrieved 2 min after tail vein-injection of VEGF-A (0.25µg/g body weight),PV (50 µmol/g body weight) or PBS, and immunoblotting for Paladin and VEGFR2. IP with isotype control IgG as negative control (IgG con). Each lane represents lysate from one mouse lung. **(d)** Representative image of co-localization (yellow) of Paladin (green) and VEGFR2 (red) in VEGF-A–treated HDMEC (50 ng/ml; 10 min), observed using super-resolution microscopy. Scale bars: 10 μm and 0.5 μm (inset). n=15. **(e)** Cell surface VEGFR2 levels detected by cell surface biotinylation, using thiol-cleavable sulfo-NHS-SS-biotin, of HDMEC transfected with *PALD1* siRNA (#1 and #2) or non-targeting control (‘c’) siRNA, followed by VEGF-A stimulation (50 ng/ml) for indicated time periods. Total lysates and streptavidin (SA) pull down, immunoblotted for VEGFR2, Paladin, and actin. ‘No biotin’, cells not treated with sulfo-NHS-SS-biotin. **(f)** Quantification of data in (e). VEGFR2 surface levels (data pooled for the indicated time points) normalized to total VEGFR2 levels and compared between control and siRNA treated HDMEC. Mean±SEM. n=4. **(g)** Internalized pool of VEGFR2 after VEGF-A treatment (50 ng/ml) of non-transfected HDMEC (NT) or HDMEC transfected with *PALD1* siRNA (KD#1 and KD#2) or non-targeting control siRNA (**c**). At indicated time points, remaining cell surface biotin was stripped and the internalized pool of VEGFR2 was collected by SA pull down. Immunoblotting of the total lysate verified *PALD1* siRNA-mediated knockdown. **(h)** Quantification of data in (g). Data were normalized to total VEGFR2 levels in the lysate after subtraction of signals in biotinylated and stripped samples. Mean±SEM, *t*-test for individual time points, normalized to cntrl siRNA sample. n=3. **(i)** Analysis of endosomal vesicles following *PALD1* knockdown. Representative images of VEGFR2 (red)/EEA1 (green) double-positive (yellow) vesicles in non-transfected, negative-control siRNA, and *PALD1* KD#2 siRNA-silenced HDMEC at 30 min of VEGF-A stimulation (50 ng/ml). DAPI in blue. **(j)** Quantification of VEGFR2-EEA1 double-positive vesicles (left) and VEGFR2 vesicles (right). Mean±SEM, two-way ANOVA. n=4.

To validate a role for Paladin in VEGFR2 signaling, Paladin expression was silenced in endothelial cells. Interestingly, siRNA-mediated knockdown of *PALD1* in HDMEC resulted in a 35-51% increase of the total basal VEGFR2 pool (*PALD1* siRNA #1 (95% CI 19-51% p-value <0.05, siRNA #2 95% CI 21-81%, p<0,001) (Suppl Figure 2a). However, the receptor was degraded at the same rate as control-treated cells after VEGF-A stimulation (Figure 2e, Suppl Figure 2b). To study the effect of the presence and absence of Paladin on the trafficking of surface VEGFR2, we treated endothelial cells, in which *PALD1* expression had been silenced and non-silenced cells, with VEGF-A for different time periods. We then used surface biotinylation to pull down VEGFR2 using streptavidin separating the cell surface-localized VEGFR2 pool from the internal pool. When normalized to total VEGFR2 levels, the amount of VEGFR2 at the cell surface in control and *PALD1* siRNA-treated cells was not significantly different after VEGF-A treatment (Figure 2e,f and Suppl Figure 2c). In a parallel analysis, we evaluated the size of the internalized VEGFR2 pool over time, after stripping the cell-surface biotin-labelled VEGFR2 pool. After 15-min treatment with VEGF-A and normalization to total VEGFR2, the internalized VEGFR2 pool in *PALD1*-silenced endothelial cells was almost twice that of the control culture (Figure 2g,h and Suppl Figure 2d). This suggested that Paladin controls the rate of VEGFR2 internalization at the early time points after VEGF-A stimulation.

Finally, we stained *PALD1* siRNA-treated cells for the presence of VEGFR2 and the early endosome marker EEA1, tracking the appearance and disappearance of VEGFR2 over time as a way to indirectly assess the movement of VEGFR2 through the early endosome compartment. A similar proportion of VEGFR2^+^ vesicles co-localized with EEA1 10 min after VEGF-A stimulation in silenced and control cells, while fewer VEGFR2^+^ vesicles co-localized with EEA1 30 min after the stimulation in *PALD1*-silenced cells than in control cells (Figure 2i,j and Suppl Figure 2e-f). This suggested that VEGFR2 moved more quickly through the early endosome compartments after VEGF-A stimulation in the absence of Paladin.

Collectively, these analyses indicated that Paladin acts as a negative regulator of VEGFR2 internalization and early endosome transport upon VEGF-A stimulation.

### Loss of Paladin leads to increased ERK1/2 phosphorylation downstream of VEGFR2

Having shown that Paladin plays a role in VEGFR2 trafficking in response to VEGF-A, we next wished to explore whether VEGFR2 signaling was affected by the loss of *PALD1*.

Accordingly, we investigated the effect of *PALD1* silencing on the phosphorylation of various molecules involved in the VEGF-A/VEGFR2 signaling in HDMEC. In keeping with a more efficient internalization of the receptor after VEGF-A stimulation of HDMEC, phosphorylation of VEGFR2 normalized to total VEGFR2 was enhanced in *PALD1* siRNA-treated cells after VEGF-A stimulation compared with non-silenced cells (Figure 3a,b). Furthermore, the phosphorylation of certain downstream targets of VEGFR2: mitogen-activated protein kinase (MAPK)3/MAPK1 (ERK1/2) and SRC were increased in silenced HDMEC (Figure 3 c,d and Suppl Figure 3a), while there was no significant difference in the phosphorylation of another downstream target AKT (Suppl Figure 3b). These observations indicated that distinct pathways downstream of VEGFR2 were hyper activated in the absence of Paladin in HDMEC.

**Figure 3.**
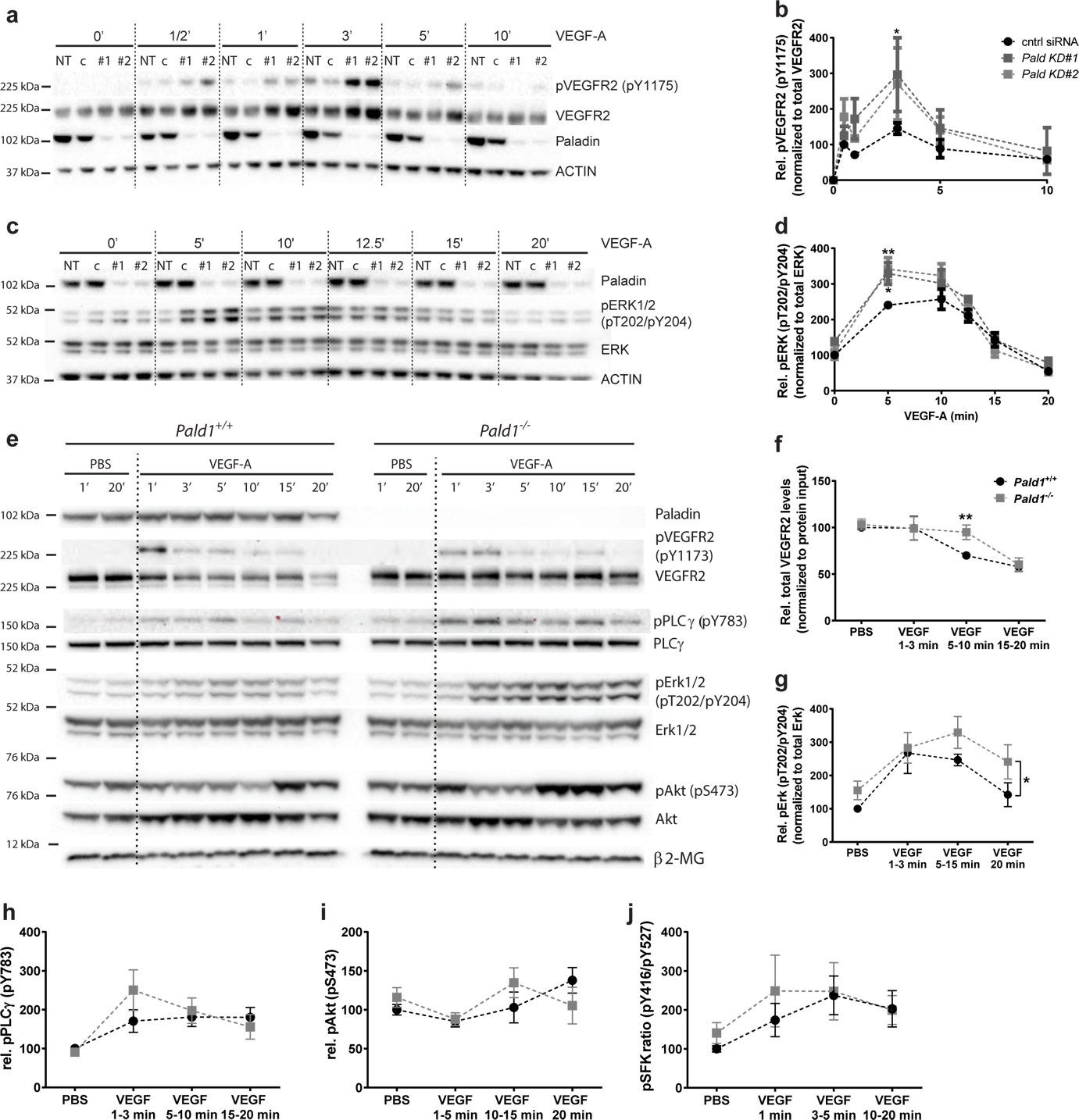
Paladin regulates VEGFR2 signaling. **(a–d)** Signaling downstream of VEGF-A/VEGFR2 assessed in HDMEC, untreated, transfected with non-targeting siRNA (NT/cntrl), or with two different *PALD1*-targeting siRNAs (KD#1 and KD#2), and treated with VEGF-A for 0-20 min. Immunoblotting of cell lysates for phosphorylated (p) VEGFR2 (pY1175), total VEGFR2, phosphorylated Erk1/2 (pT202 and pY204), and total Erk. Actin served as loading control. *PALD1* knockdown was verified by blotting for Paladin. Quantification of pVEGFR2 (normalized to total VEGFR2) (b) and pErk1/2 (normalized to total Erk1/2) (d). Mean±SEM, two-way ANOVA. n=3. **(e–j)** Immunoblotting on total heart lysates from *Pald1*^+/+^ and *Pald1*^-/-^ mice, tail-vein injected with VEGF-A or PBS for the indicated time points, for Paladin, phosphorylated and total levels of VEGFR2, phospholipase Cγ (PLCγ), Erk1/2, and Akt, and *β*2-microglobulin (*β* 2-MG, loading control) (e). Quantification of total VEGFR2 levels normalized to total loading control (f), pT202/pY204 Erk1/2 normalized to Erk1/2 (g) pY783 PLCγ normalized to PLCγ (h), pS473 Akt normalized to Akt (i), pY416 Src family kinase (SFK) normalized to pY527 SFK (j). Blot for total c-Src is shown in Suppl. Figure 3c. Mean±SEM, multiple *t*-test (Holm-Sidak for panel e), two-way ANOVA. n=4-5.

We next investigated the loss of *Pald1* on VEGFR2 signaling *in vivo*. We examined murine cardiac endothelial cells as an example of microcirculatory endothelial cells responsive to VEGF-A. Furthermore, heart endothelial cells express *Pald1* and, in contrast with the lung endothelial cells, they do not show an overt phenotype in the *Pald1*^-/-^ mouse (Wallgard, Nitzsche et al. 2012, Egana, Kaito et al. 2017, Tabula Muris, Overall et al. 2018). Accordingly, we injected VEGF-A into the tail vein of adult mice, isolated the hearts at predetermined time points and analyzed signaling proteins by western blot. Compared with the wild-type *Pald1* mouse, VEGFR2 degradation was delayed in *Pald1*^-/-^ heart lysates, which manifested as less degradation 5–10 min after VEGF-A stimulation. 15–20 min after VEGF-A stimulation *Pald1*^+/+^ VEGFR2 levels were equivalent to *Pald1*^-/-^ (Figure 3e,f). Furthermore, downstream signaling in the *Pald1*^-/-^ hearts was altered. After VEGF-A stimulation, Mapk3/Mapk1 (Erk1/2) phosphorylation was increased and prolonged in *Pald1^-/-^* mice compared with the wild-type littermates (Figure 3e,g). Other downstream mediators of VEGFR2 signaling also appeared to be aberrantly activated in *Pald1*^-/-^. However, we did not observe statistically significant differences in the level of phosphorylation of phospholipase (PLC) γ, Akt, or Src in response to VEGF-A *in vivo* in *Pald1*^-/-^ compared to wild-type littermates (Figure 3h–j and Suppl Figure 3c).

Taken together, loss of Paladin function *in vitro* and *in vivo* results in altered activation and degradation of VEGFR2, and increased Erk1/2 signaling downstream of VEGFR2.

### Endothelial hypersprouting in the *Pald1*-deficient postnatal retina results from exaggerated Erk1/2 signaling

Retinal endothelial cells proliferate and migrate in a VEGF-A/VEGFR2-dependent and highly stereotyped manner during early postnatal development (Gerhardt, Golding et al. 2003). We therefore investigated the consequence of *Pald1* gene inactivation on blood vessel development of early postnatal retinas in mice. We observed a reduced vascular outgrowth in the *Pald1*^-/-^ retina, with the most prominent effect at postnatal day (P) 5 (Figure 4a,b). Further, filopodia and tip cell numbers were higher in the *Pald1^-/-^* mouse (Figure 4c,d and Suppl Figure 4a,b). In addition, the density of the vascular front in *Pald1*^-/-^ P5 retina was greater in both the capillaries and around veins, as compared to the littermate control retina (Figure 4 e,f). These observations indicated that Paladin is a negative regulator of angiogenic sprouting. Furthermore, in agreement with an increased pErk1/2 accumulation in response to VEGF-A stimulation, as detected by immunoblotting of *Pald1*^-/-^ hearts (Figure 3e,g), the area of vascular Erk1/2 immunostaining was enhanced in the *Pald1^-/-^* retina (Figure 4g,h). Since cyclin D1 is a transcriptional target downstream of pErk1/2, we also examined the expression and localization of this protein in the retina. The analysis confirmed nuclear staining of cyclin D1 in endothelial cells and increased levels of the *Ccnd1* mRNA in *Pald1^-/-^* retina compared with the levels in littermate control retinas (Figure 4i,j). To study whether the observed phenotypes in the *Pald1^-/-^* retina could be due to increased Erk1/2 signaling, we employed an inhibitor of MAP2K1 (MEK), a Ser/Tyr/Thr kinase upstream of Erk. Notably, Erk1/2 phosphorylation in P5 retina was reduced in a time-dependent manner after a single intraperitoneal dose with the MEK inhibitor U0126 (Suppl Figure 4c). We observed that treatment with the MEK inhibitor U0126 normalized the vascular outgrowth and endothelial tip cell numbers in *Pald1^-/-^* retinas, which underscored the contribution of the exaggerated Erk1/2 signaling to the observed phenotype (Figure 4k,l). Collectively, these observations suggested that Paladin is a negative regulator of Erk1/2 signaling and endothelial sprouting and that the hypersprouting phenotype seen in *Pald1*^-/-^ retinas can be normalized by inhibition of MEK activity in the early postnatal retina in mouse.

**Figure 4.**
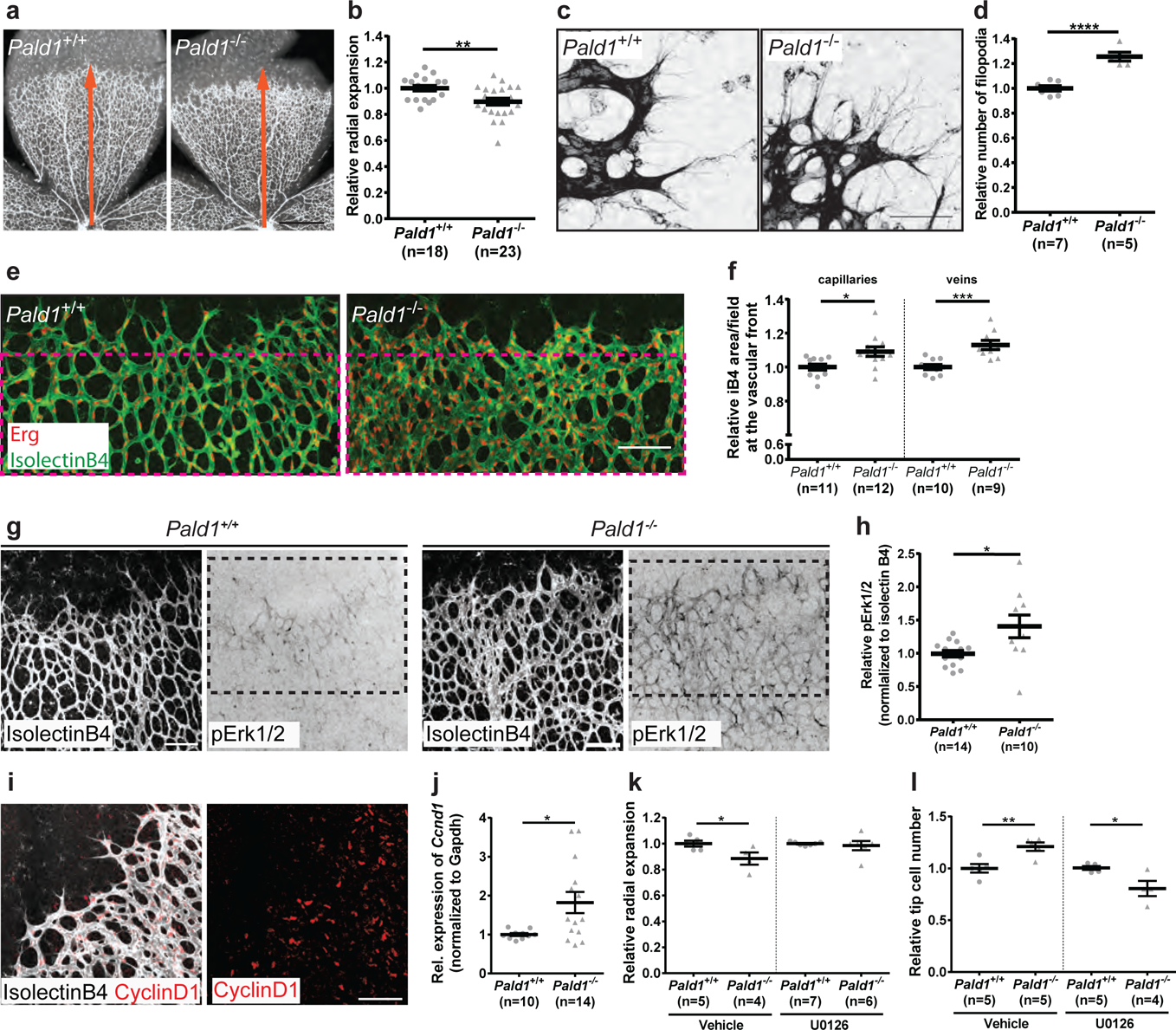
Retinal vascular phenotype in *Pald1^-/-^* mouse. **(a, b)** Delayed vascular outgrowth and hyperdense vascular front in isolectin B4-positive P5 retina from *Pald1*^-/-^ mouse compared with *Pald1*^+/+^ (a). Orange arrow indicates radial expansion of the vascular plexus in the *Pald1*^+/+^ retina. Scale bar: 1 mm. Quantification of radial expansion (b) as shown in a) normalized to wild type litter mates. Mean±SEM, unpaired *t*-test. n=14 litters, 18 *-* 23 pups per genotype. **(c,d)** Increased filopodia number in *Pald1*^-/-^ mouse retina at vascular front, visualized by isolectin B4 staining (c). Scale bar: 50 μm. Quantification of filopodia (d), normalized to wild type litter mates. Mean±SEM, unpaired *t*-test. n=3 litters, 5-7 pups per genotype. **(e,f)** P5 retina vascular front (isolectin B4, green) and endothelial nuclei (Erg, red) (e). Scale bar: 100 μm. Magenta stippled square area quantified in (f). Vascular density was determined in the capillary bed (left) and in area around veins (right). Mean±SEM, unpaired *t*-test. n=4 litters, 9-12 pups per genotype. **(g,h)** pT202/pY204 Erk1/2 immunostaining (black) in *Pald1^+/+^* and *Pald1^-/-^* P5 pups. Isolectin B4 (white) visualizes the entire vasculature. Scale bar: 100 μm. Quantification of pT202/pY204 Erk1/2 area (h) as in the black stippled square in g) (400µm from the retina rim), normalized to isolectinB4 area. Mean±SEM, unpaired *t*-test. n=5 litters, 10-14 retinas per genotype. **(i)** Representative images of cyclin D1 immunostaining in P5 retina showing nuclear cyclin D1 (red) localization in retinal vessels (isolectin B4, white). Scale bar: 100 μm. **(j)** Quantitative real-time PCR analysis of the P4-P5 retinas. *Ccnd1* transcript levels, normalized to *Gapdh*. Mean±SEM, unpaired *t*-test. n=10 *Pald1*^+/+^ and 14 *Pald1*^-/-^ pups. **(k)** Quantification of relative radial expansion in vehicle (n=2 litters, 5 *Pald1*^+/+^ and 4 *Pald1*^-/-^ pups) and MEK inhibitor (U0126)-treated pups (n=4 litters, 7 *Pald1*^+/+^ and 5 *Pald1*^-/-^ pups). MEK inhibitor/vehicle was administered twice at 12h interval at P4 and eyes collected at P5. Each dot is one mouse. Mean±SEM, one-way ANOVA. n=4–7. **(l)** Quantification of the tip cell number in vehicle-(n=3 litters, 5 *Pald1*^+/+^ and 5 *Pald1*^-/-^ pups) and MEK inhibitor (U0126)-treated pups (n=3 litters, 5 *Pald1*^+/+^ and 4 *Pald1*^-/-^ pups). MEK inhibitor/vehicle administered twice at P5 at 2-h intervals, and eyes collected 2 h after the second injection. Each dot is one mouse. Mean±SEM, one-way ANOVA.

### Absence of *Pald1* leads to increased pathological retinal angiogenesis

Pathological retinal angiogenesis is induced in hypoxia and VEGF-A is a known driver of the pathology in such diseases as wet age-related macular degeneration, where VEGF-A blockade is an important treatment (Mitchell 2011). Since we identified a role for Paladin in regulating endothelial sprouting and VEGF-A/VEGFR2 signaling, we decided to investigate the importance of *Pald1* in pathological retinal angiogenesis. We utilized an oxygen-induced retinopathy (OIR) model in mice to trigger vaso-obliteration and compensatory pathological angiogenesis (Connor, Krah et al. 2009). We used the *Pald1*^+/LacZ^ mouse to track activation of *Pald1* transcription. We observed strong endothelial LacZ expression in the retinal vasculature and in pathological blood vessels in the *Pald1*^+/LacZ^ retina following OIR (Figure 5a). Indeed, VEGF-A induced production of the Paladin protein in endothelial cells *in vitro* and in the retinal vasculature *in vivo*, as indicated by LacZ reporter expression (Figure 5b,c and Suppl Figure 5a). In addition, mice lacking *Pald1* exhibited increased vascular tuft formation at P17, but showed no difference in avascular area, compared with wild-type mice following OIR (Figure 5d-f and Suppl Figure 5b). Of note, we did not observe any differences in the vascular leakage in wild-type and *Pald1*^-/-^ after OIR, based on microsphere extravasation, or in the phosphorylation of the endothelial adherens junction protein VE-cadherin (VEC) in the vascular tufts at P17 (Suppl Figure 5c–f). Hence, expression of Paladin could be upregulated by VEGF-A and loss of Paladin specifically affected pathological vascular tuft formation in the OIR model but did not affect vascular integrity. In addition, as observed at the early developmental stage, pErk1/2 immunostaining intensity was increased in the vasculature at P15 in *Pald1*^-/-^ retinas compared to *Pald1*^+/+^ in the OIR model (Figure 5g,h).

**Figure 5.**
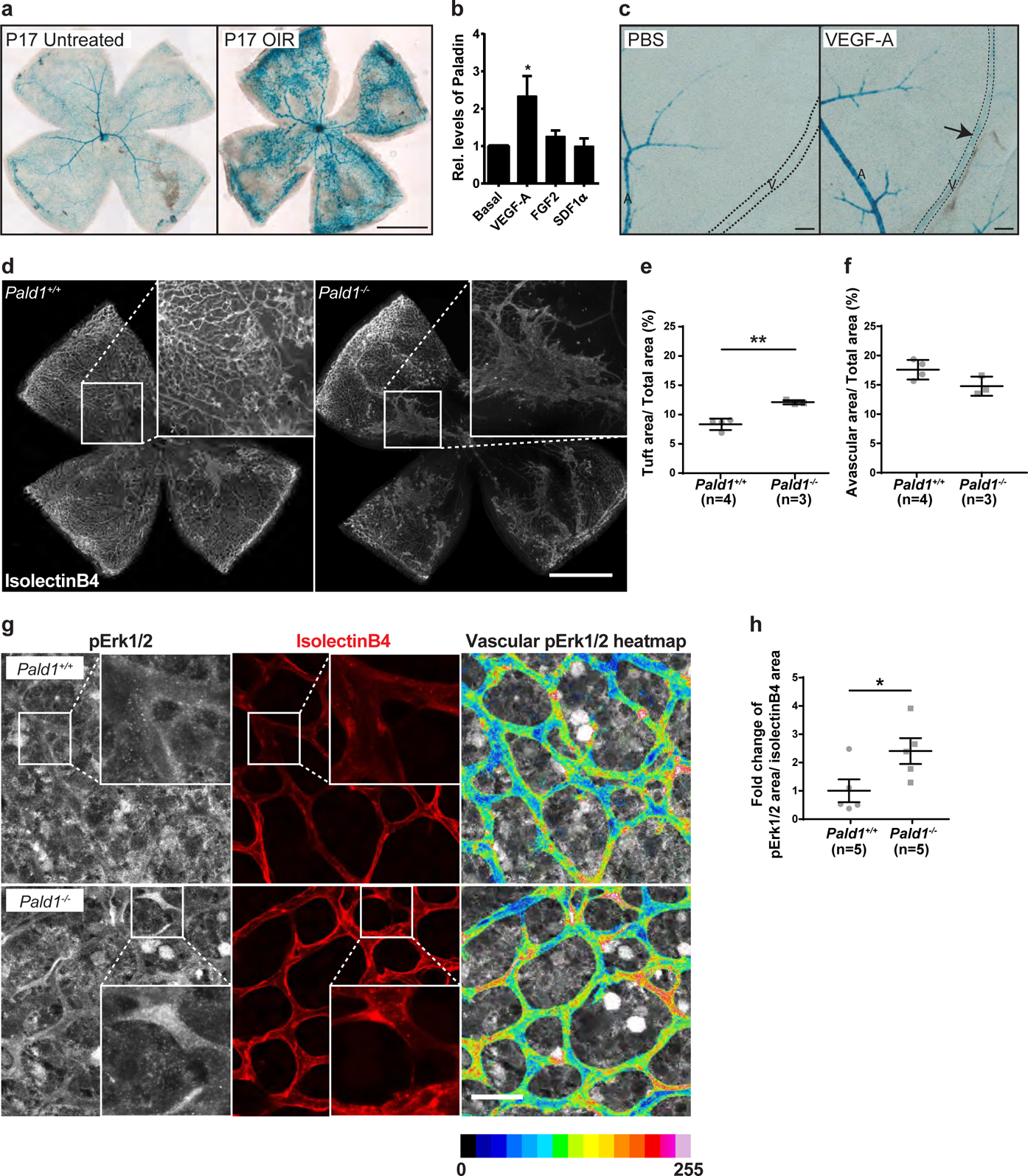
Paladin is induced by VEGF-A and regulates Erk phosphorylation in pathological angiogenesis. **(a)** Eyes from *Pald1*^+/LacZ^ mice collected at P17 from untreated animals or animals with oxygen-induced retinopathy (OIR). *Pald1*-promoter driven LacZ expression and X-gal staining generated signals in capillaries, veins and arteries in the OIR retina at P17 compared to the normoxia control with predominantly arterial LacZ expression. Scale bar: 1 mm. **(b)** Paladin levels in primary human umbilical vein endothelial cells (HUVEC) untreated or treated for 24 h with VEGF-A (50 ng/ml), FGF2 (50 ng/ml), or SDF1*α* (30 ng/ml), and immunoblotting for Paladin and *β*-actin (loading control). Mean±SEM, one way-ANOVA. n=5. **(c)** Eyes from adult *Pald1*^LacZ/+^ mice collected at 72 h after single-bolus intravitreal injection of 1 μg VEGF-A, and PBS in the contralateral eye, followed by X-gal staining. Arrow indicates *Pald1* promoter activity in veins specifically after VEGF-A treatment. A, artery; V, vein. Scale bar: 100 μm. **(d-f)** Representative images of isolectin B4-stained P17 retinas from OIR-challenged *Pald1*^+/+^ and *Pald1*^-/-^ mice (detailed view in the insets). Scale bar: 1 mm. Quantification of neovascular tuft area (e) and avascular area (f). Each dot represents the mean of both retinas per mouse. Mean±SEM, unpaired *t*-test. n=3 litters, 4 *Pald1*^+/+^ and 3 *Pald1*^-/-^ pups. **(g)** Representative images of retina vasculature immunostained for isolectin B4 and pT202/pY204 Erk1/2 (pErk1/2). pErk1/2 staining within the vessels is also visualized using a 16-color heatmap to display staining intensity. Scale bar: 30 µm. **(h)** Quantification of ppErk1/2 immunostaining as shown in g) within isolectin B4-positive vessels, as fold-change of pErk1/2 stained area. Each dot represents the mean of both retinas per mouse. Mean±SEM, unpaired *t*-test. n=3 litters, 5 *Pald1*^+/+^ and 5 *Pald1*^-/-^ pups.

Taken together, Paladin is upregulated by VEGF-A, which is the main driver of pathological retinal angiogenesis in mouse and human, and in the OIR model Paladin functions as a negative regulator of pathological angiogenesis.

## DISCUSSION

In this study, we explored the signaling role of Paladin. We show here for the first time that Paladin binds to and dephosphorylates phosphoinositides. Paladin was localized to Rab4, −7, and −11, and Golgi membranes, and was thus positioned to act as a regulator of endosomal phosphoinositide content and, thereby, endosomal trafficking. Furthermore, loss of *Pald1* expression led to enhanced VEGFR2 internalization after VEGF-A stimulation, rapid trafficking of VEGFR2 through early endosomes, and increased pERK1/2 levels *in vitro* and *in vivo*. Phenotypically, *Pald1* deficiency resulted in retinal endothelial hypersprouting, which could be normalized by inhibition of MEK, and enhanced pathological retinal angiogenesis. Our data suggest that Paladin is a part of a VEGF-driven negative feedback loop in retinal angiogenesis where VEGF-A upregulates Paladin which acts to dampen VEGFR2 driven signaling and endothelial sprouting. Through inhibition of Paladin one could potentially activate VEGF-A/VEGFR2 signaling in e.g. situations of insufficient angiogenesis.

Despite the lack of published experimental evidence, Paladin had been postulated to be a catalytically inactive pseudophosphatase (Huang, Hancock et al. 2009, Kharitidi, Manteghi et al. 2014, Reiterer, Eyers et al. 2014). However, more recently, Alonso and Pulido suggested that Paladin is a Cys-based phosphatase, which forms its own subclass (IV). Whereas all the neighboring phosphatase subclasses dephosphorylate phosphoinositides, they proposed, based on structural similarity, that Paladin might possess inositol phosphatase activity (Alonso and Pulido 2015). Indeed, we have demonstrated in this study that Paladin, while also binding monophosphoinositides and PI(3,5)P_2_, preferentially dephosphorylates PI(4,5)P_2_ lipids, but not other phosphoinositides or inositol phosphates. As has been shown for PTEN, the lipid-binding specificity might serve to localize Paladin to particular trafficking vesicles (Naguib, Bencze et al. 2015). Based on the presented findings, we propose that Paladin controls endosomal vesicular trafficking in endothelial cells by modifying the lipid membrane of intracellular vesicles. In line with this hypothesis, Paladin was localized to endosomal vesicles, co-localized with Rab4, −7, and −11 positive vesicles, and also appeared in close association with the Golgi membranes. As one possibility, based on the substrate preference and combined with the subcellular localization of Paladin, the protein might bind PI(3,5)P_2_ in late endosomes, and control the levels of PI(4,5)P_2_ in sorting or recycling endosomes (Tan et al JCS 2015). However, further studies are needed to formally verify this.

Paladin appears to be a PI(4,5)P_2_ phosphatase with a specific biological function, as has also been observed for other PI(4,5)P_2_ phosphatases, which have similar substrate specificity but play distinct biological roles. For example, syntaptojanins, endocytic PI(4,5)P2 phosphatases, appear to play a role in neurological functions such as synaptic vesicle recycling and loss of function leads to neurological disorders and seizures in both mice and humans (Dyment, Smith et al. 2015, Nakatsu, Messa et al. 2015), while OCRL1, a PI(5)P phosphatase, is mainly involved in the development of eye, brain and kidney (Attree, Olivos et al. 1992). We have previously shown that Paladin is essential for normal lung development as female mice exhibit an emphysema-like phenotype and show increased turnover of endothelial cells in the lung (Egana, Kaito et al. 2017). In addition, we have now demonstrated that Paladin is involved in retinal vascular development and pathology. Further research will be needed to both dissect substrate specificity at subcellular resolution and also to connect it with physiological roles at the organismal level for PI(4,5)P2 phosphatases to understand, ultimately, how enzymes with the same substrate specificity act in concert to regulate potent signaling lipids.

Paladin appears to play a unique VEGF-A–regulated role by affecting VEGFR2 trafficking and signaling. Indeed, we observed that following VEGF-A stimulation, VEGFR2 was more rapidly internalized, with enhanced ERK1/2 phosphorylation, in *PALD1* silenced cells than in the control cells. Disruption of VEGFR2 trafficking has also been shown to occur in Synectin deficiency. Synectin null mice exhibit altered VEGFR2 signaling and insufficient arterial formation as a consequence of prolonged trafficking through early endosomes which allows for increased VEGFR2 dephosphorylation at pTyr1173 by tyrosine-protein phosphatase non-receptor type 1 (PTP1b) and reduced pERK1/2 (Lanahan, 2010). Hence, in the absence of *Pald1*, the rapid internalization of VEGFR2 may allow its escape from dephosphorylation by PTP1b and, thereby, increased pERK1/2 levels. Alternatively, VEGFR2 might recycle back to the plasma membrane and initiate another round of phosphorylation in the absence of Paladin. In line with this, we also observed increased pTyr1173 phosphorylation in HDMEC but did not compare it to the levels of other VEGFR2 phosphosites. Future studies could address the possible time- and phospho-site specific effect of Paladin loss-of-function and the association of specific tyrosine phosphatases to VEGFR2 to address the similarities and differences between Paladin and Synectin loss of function.

Retinal angiogenic sprouting is driven by VEGF-A/VEGFR2 signaling (Gerhardt, Golding et al. 2003), which is regulated by receptor internalization and subcellular localization (Gaengel and Betsholtz 2013, Nakayama, Nakayama et al. 2013, Simons, Gordon et al. 2016). We observed hyperphosphorylation of Erk1/2 as well as hypersprouting in the *Pald1* knockout retina and in pathological vessels in the OIR. This is in line with the observation that Erk1/2 is one important downstream mediator of angiogenic sprouting; it was recently shown that Erk1/2 signaling is both necessary and sufficient for endothelial tip cell sprouting in zebrafish (Shin, Beane et al. 2016). However, signaling alterations in the *Pald1*^-/-^ retina are likely more complex, as defective endosomal trafficking not only affects VEGFR2 but most likely also other signaling molecules. Nevertheless, we observed normalization of the increased endothelial tip cell numbers and vascular outgrowth defects in the *Pald1* knockout retina upon MEK inhibitor treatment, underscoring the role of Erk1/2 activation as an important part of the signaling defects in *Pald1*-deficient endothelial cells. Recently, *PALD1* has been genetically associated with Moyamoya disease in two families. Moyamoya disease is caused by the occlusion of the carotid artery and its branches, causing a characteristic pronounced collateral vessel formation and stroke in the central nervous system (Grangeon, Guey et al. 2019). A potential causal link between the hyperactive endothelial signaling observed in the *Pald1*^-/-^ mouse and excessive collateral formation in patients with Moyamoya disease should be studied further.

Based on the evidence presented herein, we propose that Paladin is a critical regulator of VEGFR2 endosomal trafficking. Potentially, Paladin might also regulate the activity of other membrane proteins and receptors. Indeed, Paladin interacts with TLR9, an entirely endosomal signaling receptor, and reduced Paladin expression leads to blunted TLR9 signaling (Li, Wang et al. 2011). Paladin has also been identified as a negative regulator of insulin receptor signaling, and *PALD1* deficiency leads to increased insulin receptor levels and increased AKT downstream signaling (Huang, Hancock et al. 2009). Considering the diversity of receptors affected by Paladin, its activity as a phosphoinositide phosphatase may be a common denominator of such general role in membrane protein trafficking and signaling.

In conclusion, we have demonstrated that Paladin is a VEGF-A–inducible lipid phosphatase that regulates endothelial sprouting and VEGFR2 trafficking and signaling, likely exerting its effects by controlling the level of phosphoinositides in the sorting/recycling endosomal compartment. Therapeutic targeting of VEGFR2 signaling indirectly via the inhibition of PI phosphatases such as Paladin might be a way to modify of VEGFR2 signaling that is not achievable by targeting the receptor directly. For example inhibition of Paladin, or other lipid phosphatases, could be used to enhance VEGFR2 signaling to promote angiogenic sprouting in situation of insufficient angiogenesis.

## ACKNOWLEDGEMENTS

We thank Kurt Ballmer-Hofer for kindly sharing canine VEGF-A_164_. We thank Elisabetta Dejana for the kind gift of pY685 VE-cadherin antibodies. We thank Carina Hellberg (deceased), Katie Bentley, and Andrew Philippides for valuable discussion and input. High-resolution SIM images, bright field, and fluorescent images were acquired at the SciLife Lab BioVis Platform at The Rudbeck Laboratory, Uppsala University, Sweden. This work was supported by the Swedish Cancer Foundation, Beijer Foundation, Åke Wiberg’s Foundation, Magnus Bergwall’s Foundation, and Swedish Research Council (E.E., L.C.-W.); Knut and Alice Wallenberg Foundation (L.C.-W.); and Gustav Adolf Johansson Foundation (J.L., I.E.).

## AUTHOR CONTRIBUTIONS

Experimental design, generation, and analysis of data: A.N., R.P. C.T., T.N., R.O.S., E.E., J.L., F.P.R., I.E., S.J., L.C.-W., M. H. Manuscript figure assembly: A.N., R.P., C.T. Reagents: P.B. Supervision: M.H. Manuscript writing: A.N., R.P., C.T., P.B., L.C.-W., M.H.

## COMPETING INTERESTS

The authors declare no competing interests.

## FIGURE SUPPLEMENTS

Supplementary Figure 1. Paladin lacks protein phosphatase activity *in vitro*. **(a)** Amino acid sequence alignment of the third (amino acids: mouse, 312-–322; human, 309–319) and fourth (amino acids: mouse, 661–671; human, 658–668) putative phosphatase domains of Paladin with the consensus sequence of catalytic domain motif 9 of cysteine-based protein tyrosine phosphatases revealed Ser instead of His residue in front of the Cys in the phosphatase domains. By contrast, the phosphatase domain of vascular endothelial PTP (VE-PTP) contains the complete PTP motif. **(b)** Screening of the phosphatase activity towards phosphoinositides and inositol phosphates of immunoprecipitated wild-type full-length Paladin and phosphatase-dead (C/S) mutant variant using an *in vitro* colorimetric molybdate dye assay. Commercial SHIP2 enzyme reaction buffer was used (Echelon, USA). Mean±SEM, n=3 replicates, one experiment. **(c)** Immunoprecipitated full-length wild-type Paladin or its phosphatase-dead C/S variant expressed in HEK293 cells were analysed in an *in vitro* radioactive phosphatase assay using phosphorylated PKC-optimal peptide containing phosphoserine and phosphothreonine residues as a substrate. Immunoprecipitates of cells transfected with an empty vector served as a negative control. Data were normalized to ^32^P input. Mean±SEM. n=7 for full-length Paladin and n=3 for C/S Paladin variant (individual experiments). **(d)** Representative image of co-localization of Paladin (green) and Vascular Endothelial Cadherin (VEC, red) shown by super-resolution microscopy of VEGF-A–treated HDMEC (50 ng/ml; 10 min). Scale bar: 10 μm and 2 μm (inset). n=5. **(e)** Paladin binds mono- and diphosphoinositides *in vitro*. Binding capacity of full-length GST-tagged Paladin to immobilized phosphoinositides was assessed using a PIP-array membrane. The amount of respective phosphoinositide is listed in pmol. **(f)** Confocal representative image of HDMEC stained Paladin (green) and the Golgi apparatus marker giantin (red). Cells were treated with VEGF-A (50 ng/ml) for 10 min. Scale bar: 5 μm. n=4. **(g)** Confocal representative image of VEGF-A treated HDMEC (50 ng/ml; 10 min) immunostained for the presence of Paladin (green) and the lysosomal vesicle marker Rab7 (red), after transduction with Rab7-cherry baculovirus. Note the co-localization of Paladin and Rab7 (yellow). Scale bars: 10 μm and 2 μm (inset). n=2. **(h)** Confocal representative image of HDMEC stained for the presence of Paladin (green) and Early Endosome Antigen 1 (EEA1, red). Cells were treated with VEGF-A (50 ng/ml) for 10 min. Scale bar: 10 μm and 2 μm (inset). n=4.

Supplementary Figure 2. Paladin regulates VEGFR2 endosomal trafficking, total and internalized VEGFR2 levels after VEGF-A stimulation of HDMEC. (a) Total VEGFR2 was quantified in HDMEC treated with control siRNA (cntrl) or siRNA targeting *PALD1* (KD#1 or KD#2) for 72h. Mean±SEM, one-way ANOVA. n=13. (b) Total VEGFR2 was quantified in HDMEC treated with control siRNA (‘c’) or siRNA targeting *PALD1* (#1 or #2), as shown in Figure 2e. Time dependent degradation was observed after VEGF stimulation in both control and *PALD1* siRNA treated cells. (c) Quantification of VEGFR2 surface levels in *PALD1*#KD1 cells from four independent experiments, as shown in Figure 2e. VEGFR2 surface levels (data pooled for the indicated time points) were normalized to total VEGFR2 levels in the lysate and compared between control and siRNA treated HDMEC. Mean±SEM. n=4. (d) Quantitative analysis of internalized VEGFR2, data from three individual experiments for *PALD1*#KD1, as shown in Figure 2g. Data were normalized to total VEGFR2 levels in the lysate after subtraction of signals in biotinylated and stripped samples. Mean±SEM, *t*-test for individual time points, normalized to cntrl siRNA sample. n=3. (e) Representative images of VEGFR2 (red)/EEA1 (green) double-positive vesicles (yellow) in non-transfected, negative-control siRNA, and *PALD1* KD#1 siRNA-silenced HDMEC after 30 min of VEGF-A stimulation (50 ng/ml). (f) Quantification of VEGFR2-EEA1 double-positive vesicles (left) and VEGFR2 vesicles (right). n=4. Mean±SEM, two-way ANOVA.

Supplementary Figure 3. **VEGF-A/VEGFR2 downstream signaling *in vitro* and *in vivo*.**. **(a,b)** HDMEC, untransfected (NT) or transfected with non-targeting (c/cntrl) or *PALD1*-targeting siRNA (KD #1, KD #2), were treated with VEGF-A for the indicated time periods. Immunoblots of cell lysates to determine protein levels of ACTIN, Paladin, pTyr416 and pTyr527 of Src family kinases (SFK) and total Src (a), pSer473 AKT, and total AKT (b),, are shown. Mean±SEM, two-way ANOVA. n=3. **(c)** *Pald1*^+/+^ and *Pald1*^-/-^ mice were tail-vein injected with VEGF-A or PBS for the indicated time periods. Heart lysates were blotted to determine the phosphorylated and total levels of SFK. Quantification in Figure 3j. n=5 individual experiments.

Supplementary Figure 4. Loss of *Pald1* leads to hypersprouting of the retinal vasculature. **(a)** Number of filopodia per 100-μm vascular front in the P5 retina stained with isolectin B4. Mean±SEM, unpaired t-test. n=3 litters, n=5-7 pups per genotype as indicated in graph. **(b)** Nuclei of tip cells, defined as the front cells extending filopodia, at the vascular front of the P5 retina stained for endothelial cells (isolectin B4) and endothelial nuclei (Erg). Data for each litter were normalized to that of *Pald1*^+/+^ littermates. Mean±SEM, unpaired *t*-test. n=4 litters, n=7-9 pups per genotype as indicated in graph. **(c)** Western blot analysis of pT202/pY204 Erk1/2 levels in the P5 retina following a single intraperitoneal injection of MEK inhibitor U0126 (5 mg/kg), analyzed after the indicated time periods.

Supplementary Figure 5. Paladin does not affect leakage or VE-cadherin phosphorylation in pathological retinal angiogenesis. **(a)** Immunoblotting for paladin in HUVEC, untreated or treated for 24 h with VEGF-A (50 ng/ml), FGF2 (50 ng/ml), or SDF1*α* (30 ng/ml), actin served as loading control. Quantification in Figure 5b. n=5. **(b)** Representative images of isolectinB4 stained retina of P15 mouse during OIR development, showing similar levels of vessel dropout between the genotypes caused by hyperoxic conditions at P7–P12. Scale bar: 500 μm. n=3 retinas per genotype **(c)** Representative images of microsphere extravasation from neovascular tufts following intravenous injection of 25-nm fluorescent microspheres into mice that had been subjected to OIR. White arrows emphasize the extravascular accumulation of microspheres. Scale bar: 25 μm. n=3 litters, 4 pups per genotype. **(d)** Quantification of microsphere extravasation in the *Pald1*^+/+^ and *Pald1*^-/-^ retinas at P17 during OIR, as shown in c. Mean±SEM, unpaired *t*-test. n=3 litters, 4 pups per genotype. **(e)** Representative images of isolectin B4, Vascular endothelial cadherin (VEC), and pY685 VEC immunostaining in the P17 retina of mice subjected to OIR. Scale bar: 50 μm. n=3 litters, 5-7 pups per genotype. **(f)** Quantification of phY685 VEC in the *Pald1*^+/+^ and *Pald1*^-/-^ retina at P17 during OIR, as shown in e. Mean±SEM, unpaired *t*-test. n=3 litters, 5-7 pups per genotype.

## ONLINE METHODS

### Mice

C57BL/6 mice with constitutive deletion of *Pald1* (Exon 1-18 replaced by a LacZ reporter cassette) have been generated^1^ and backcrossed for at least 10 generations. *Pald1^+/-^* intercrosses were performed to generate homozygous and heterozygous littermates. All animal experiments were performed in compliance with the relevant laws and institutional guidelines and were approved by the Uppsala University board of animal experimentation.

MEK inhibitor U0126 (V1121, Promega) was injected intraperitoneally (5 mg/kg). For the short treatment, pups were injected twice at P5 with a two hour interval and eyes were collected two hours after the last injection. For the long treatment, pups were injected twice with a 12 h interval at P4 and retinas were collected for analyses at P5. As a vehicle, 40% DMSO in sterile 1xPBS was used.

### Statistical analysis

Statistical analysis of two data sets was done by unpaired student’s t-test (normal distribution of samples was verified) and of three data sets or more was done by one-way ANOVA GraphPad Prism6 and Prism7. Statistical significance is indicated as follows: * p ≤ 0.05, ** p ≤ 0.01, *** p ≤ 0.001.

### Cell culture and reagents

HUVECs (ScienCell Research Laboratories) and HDMECs (PromoCell) were cultured in cell culture dishes coated with 1% gelatine using endothelial cell medium MV2 (PromoCell) with all supplements (5% FCS, 5 ng/ml hEGF, 0.5 ng/ml VEGF, 20 ng/ml R3 IGF, 1 µg/µl ascorbic acid, 10 ng/ml bFGF and 0.2 µg/µl hydrocortisone) at 37°C and 5% CO2. Cells between four to six passages were used.

Cells were treated with the following reagents: 50 ng/ml mVEGF-A164 (Peprotech), 50 ng/ml hVEGF-A165 (Peprotech), 50 ng/ml rh FGF2 (RD systems), 30 ng/ml rh SDF1α (ImmunoTools), 100 µM peroxyvanadate. HDMECs were starved for 2-6h or overnight in 0.1% FBS prior to growth factor stimulations.

To knock-down *PALD1* mRNA, semi-confluent HDMECs were transfected with siRNAs targeting *PALD1* (s25894, s25895, Ambion), or non-targeting siRNA (Stealth RNAi negative control, medium GC, Thermo Fisher) using RNAi Max (Invitrogen) according to manufacturer’s instructions and cells were used for experiments 72 hours later.

### Phosphatase assay

HEK293 cells were transfected with plasmids (pcDNA3.1 or pLenti7.3-V5 backbone) encoding V5-tagged human full-length paladin or mutant paladin (see Suppl figure 1a) using Lipofectamine 2000 (Invitrogen). As controls wild type PTEN (28298 by Addgene), phosphatase-dead mutant PTEN C124S (28300 by Addgene) or β-galactosidase (pLenti7.3/V5-GW/lacZ by Invitrogen) were used. Cells were washed twice with 1xTBS and lysed (0.5% Triton X-100, 0.5% sodium deoxycholate, 150 mM NaCl, 20 mM Tris, pH 7.4, 1x protease inhibitor cocktail [Roche] or 20 mM HEPES, 150 mM NaCl, 1% NP40, 1x protease inhibitor cocktail [Roche]) and immunoprecipitation was performed with antibodies targeting the V5-tag of paladin constructs (Invitrogen) or FLAG-tag of PTEN constructs (F3165, Sigma). As a positive control in protein phosphatase assays endogenous TC-PTP was immunoprecipitated (6F3 clone, MediMabs). After 2h at 4°C, lysates were incubated with Protein-G sepharose beads (GE Healthcare) for 45 min at 4°C. Subsequently, beads were washed twice with lysis buffer and once with assay buffer (25 mM Tris-HCl, 140 mM NaCl, 2.7 mM KCl, 10 mM DTT or SHIP2 reaction buffer [Echelon]) and resuspended in 100 µl (for triplicates) or 65 µl (for duplicates) of assay buffer (one 10-cm dish of HEK293 cells per triplicate or two duplicates).

Phosphoinositide phosphates (Echelon, diC8) and inositol phosphates (Echelon, IP6 by Merck) were suspended in assay buffer at 3000 pmol/well. The protein to be tested (30 µl of immunocomplexes) was added and the reaction was stopped after 20-90 min depending on phosphoinositide to be tested by adding an equal volume of molybdate dye solution (V2471 Promega) and after 15 min incubation at RT absorbance at 600 nm was measured. Released phosphate was calculated by comparison to the amount of free phosphate in positive control (3000 pmol of K2PO4). The colorimetric assay was performed in 96-well half area plates (Costar # 3690).

Phosphopeptide phosphatase activity was assessed by the radioactive assay using src-optimal peptide and PKC-optimal peptide as previously described^2^.

### PIP-array

PIP-Array membrane (Echelon) was blocked with 3% fatty-acid free BSA (Roche) in 1x PBS, 0.1% Tween (1x PBST) and incubated at 4°C overnight with 500 ng/ml of recombinant GST-tagged paladin protein (Abnova). After washing with 1xPBST, membrane was incubated at 4°C overnight with paladin antibody (HPA17343, Atlas antibodies). After secondary antibody incubation membrane was developed using enhanced chemiluminesence (GE Healthcare) with ChemiDoc MP Imaging System (Biorad Laboratories).

### Intracellular localization of Paladin

For visualization of PI(4)P and Paladin in HDMECs, cells were serum-starved for 2 hours and stimulated for 15 min. with mVEGF-A164, or left unstimulated. Cells were fixed and stained as described for visualization of the Golgi compartment^3^.

For visualization of Paladin, VEGFR2 and Rab-GTPases, HDMECs were infected with baculoviruses encoding Rab4-cherry, Rab7-cherry and Rab11-cherry^4^. Confluent HDMEC seeded on gelatine-coated 8-well chamber slides (BD Falcon) were infected with baculovirus (10% of cell culture medium) for 8 hours. The following day cells were starved overnight, stimulated for indicated time with VEGF-A and fixed in 2% paraformaldehyde (PFA) for 10 min on ice.

For immunostainings, cells were stimulated as described, fixed with 4% PFA or with cold methanol for 10-15 minutes, permeabilized if needed with 0.2% Triton-X-100 for 10 minutes and blocked in 0.2% Tween 20/3% BSA/5% FCS/0.05% Sodium Deoxycholate in PBS, or in 3% BSA-1xPBS and incubated with primary antibodies overnight at 4°C. After washing samples were incubated with fluorophore conjugated secondary antibodies (Jackson Immunoresearch) and Hoechst 33342 to visualize nuclei. Following antibodies were used: Anti-EEA1 (1:200, BD Bioscience, 610457), anti-Giantin (1:500, Alexis, AXL-804-600, or 1:100 Abcam ab24586), anti-paladin (1:50, Atlas, HPA015696), anti-PI(4)P (1:200, Z-P004 Echelon), anti-VE-Cadherin (1:200 R&D System, AF1002) and anti-VEGFR2 (1:100 R&D Systems, AF357). All samples were mounted using Fluoromount-G (SouthernBiotech) or ProLong Gold (Invitrogen).

Cells were imaged with Zeiss LSM700 or Leica SP8 confocal microscopes. Image acquisition was done with 40x and 63x objectives. Super-resolution images were acquired with Zeiss LSM710 SIM. Images were processed and quantified with ImageJ software (NIH) or Cell Profiler (Broad Institute)^5^.

### Immunoprecipitation and western blotting

Cells were washed once with cold 1xPBS and lysed in cell lysis buffer (0.02 M HEPES pH 7.5, 0.15 M NaCl, 1% [w/v] NP 40, 1 mM Na3VO4, in PBS and 1x Protease Inhibitor Cocktail (Roche)).

For *in vivo* signalling study, dog VEGF-A165 (5 µg/20 g body weight) was administrated via the tail vein and mice were sacrificed after 1-20 min circulation time, lung and heart were removed immediately and snap frozen. Control mice received an equal volume of PBS. Snap frozen tissue was lysed in 1% NP-40, 1% sodium deoxycholate, 0.01 M NaPi, 150 mM NaCl, 2 mM EDTA, 1 mM Na3VO4, 1x Protease Inhibitor Cocktail (Roche), or in 20 mM HEPES, 150 mM NaCl, 1% NP-40 with 2 mM Na3VO4 and 2.5x Protease Inhibitor Cocktail (Roche), homogenized with Tissue Tearor (BioSpec Products) and sonicated six to eight times for 5 sec at 200 W (Bioruptor, diagenode). After one-hour incubation at 4°C, tissue lysates were centrifuged at 21’100 g for 20 min. Protein concentration was measured with the BCA protein detection kit (Thermo Fisher Scientific).

For immunoprecipitation, lysates were pre-cleared for 2 h at 4°C with unspecific goat IgG (Jackson Immuno Research) and Protein-G sepharose 4 Fast Flow beads (GE Healthcare) and incubated overnight at 4°C with goat anti-mouse VEGFR2 (R&D, AF644) or goat anti-human VEGFR2 (R&D, AF357). The lysates were incubated with Protein-G sepharose beads for 1 h at 4°C and subsequently the beads were washed five times with lysis buffer and denatured in 2x sample buffer (LifeTechnologies) at 95°C for 5 min.

Proteins were separated on a 4-12% BisTris polyacrylamide gel (Novex by Life Technologies) and transferred to an Immobilon-P PVDF membrane (Millipore) using the Criterion Blotter system (BioRad). The membrane was blocked with 5% skimmed milk in TBS 0.1% Tween, or with 5% BSA in TBS 0.1% Tween for anti-phospho antibodies and incubated overnight at 4°C. Following antibodies were used. Rabbit anti-paladin (1:1000; Atlas Antibodies, HPA017343), rabbit anti-phospho-VEGFR2 pY1175 (1:1000, Cell Signaling, 2478), rabbit anti-VEGFR2 (1:1000, Cell Signaling, 2479), rabbit anti-phospho-PLCγ pY783 (1:1000, Invitrogen, 44-696G), rabbit anti-PLCγ (1:1000, Cell Signaling, 2822), rabbit anti-phospho-Erk1/2 pThr202/pTyr204 (1:1000, Cell Signaling, 4377), rabbit anti-Erk1/2 (1:1000, Cell Signaling, 9102), rabbit anti-phospho-Akt pSer473 (1:1000, Cell Signaling, 4060), rabbit anti-Akt (1:1000, Cell Signaling, 9272), rabbit anti-phospho-Src pTyr416 (1:1000, Cell Signaling, 6349), rabbit anti-phospho-Src pTyr527 (1:1000, Cell Signaling, 2105), rabbit anti-Src (1:1000 Cell Signaling, 2123) and or goat anti-actin (1:1000, Santa Cruz, sc1615). Membranes were washed in TBS 0.1% Tween and incubated with horseradish peroxidase (HRP) conjugated secondary anti-rabbit (1:10’000, GE Healthcare) or anti-goat antibodies (1:10’000, Invitrogen), respectively. Membranes were washed in TBS 0.1% Tween and developed using ECL prime (GE Healthcare). Luminescence signal was detected by the ChemiDoc MP system (BioRad) and densitometry performed using Image Lab software (BioRad).

### Surface biotinylation assay

For assessment of surface-bound VEGFR2 levels after VEGF-A stimulation, siRNA transfected HDMECs were starved for two hours in basic endothelial cell medium (PromoCell) with only 0.1% FBS and stimulated with recombinant VEGF-A164 (50 ng/ml) for indicated time points. Cells were washed twice with cold 1xDPBS (containing Mg^2+^ and Ca^2+^) and biotinylated with 0.5 mg/ml EZ-Link Sulfo-NHS-Biotin (Thermo Scientific) in DPBS at 4°C for 45 minutes with gentle shaking. The reaction was stopped by washing twice with cold DPBS and incubation with cold 100 mM glycine in DPBS for 10 minutes on ice. Subsequently, the cells were washed and lysed in modified RIPA buffer (20 mM HEPES, 150 mM NaCl, 1% NP-40) with protease (Roche) and phosphatase inhibitors (1 mM Na3VO4).

For assessment of the VEGFR2 internalized pool after VEGF-A stimulation, biotinylation of cells surface receptors was performed prior to VEGF-A stimulation as described above. Cells were stimulated with VEGF-A164 (50 ng/ml) for indicated time points. Cells were washed in cold 1x DPBS and cell-surface biotin was cleaved off by incubating the cells on ice with 100 mM of membrane impermeable reducing agent MESNA (2-mercaptoethane sulfonic acid) (Sigma) in stripping buffer (50 mM Tris, pH 8.6, 150 mM NaCl, 1mM EDTA, 0.2% BSA (pH 8.6) for 3x 15 minutes. After washing cells were lysed with RIPA buffer as described above.

Equal amounts of protein lysates were immunoprecipitated with streptavidin sepharose beads (GE Healthcare) overnight at 4°C after which beads were washed extensively with RIPA buffer and suspended in 2x NuPAGE LDS Sample Buffer (Invitrogen) with NuPAGE Sample Reducing Agent (Invitrogen). Protein separation and western blotting was performed as described above.

### Retina preparation, whole mount staining and imaging

Eyes were harvested and either fixed in 4% PFA for 10 min at RT, dissected and post-fixed in ice-cold methanol for at least 2 h (pErk; for filopodia analysis), fixed with 2% PFA for 5 h (Erg), 4% PFA for 1h at RT (CyclinD1) or fixed with 1% PFA, 0.1% triethanolamine, 0.1% TritonX-100, 0.1% NP-40 for 2 hours at RT, retinas dissected and post-fixed in ice-cold methanol for 2 h (VEC). After rehydration retinas were permeabilized and blocked (0.1-0.5% Triton-X100, 0.05% sodium deoxycholate, 1% BSA, 2% FBS, 0.02% sodium azide in PBS, or 0.3% Triton-X100, 3% FBS, 3% donkey serum) for 1-2 h at RT and stained overnight at 4°C using the following antibodies: rabbit anti-Erg (1:300, Abcam, ab92513), rabbit anti-cyclinD1 (1:50, Thermo Scientific RM-9104), rabbit anti-pERK1/2 (1:100, Cell Signaling #9101), rat anti-VEC (1:100, BD 555298) and rabbit anti-pY685 VEC (1:50, kind gift from Elisabetta Dejana^3^). After washing, retinas were incubated with appropriate fluorophore-coupled secondary antibodies and fluorophore conjugated isolectinB4 (Jackson Immunoresearch) or washed in PBlec (1% Triton X-100, 0.1 mM CaCl2, 0.1 mM MgCl2, 0.1 mM MnCl2 in PBS at pH 6.8) for at least 1 hour at RT and stained with biotinylated isolectinB4 (Sigma) overnight at 4°C and incubated with Streptavidin-Alexa 488 (Invitrogen). Retinas were flat-mounted in Fluoromount-G (Southern Biotech), or in ProLong Gold (Invitrogen).

Images were acquired with LSM700, AxioImager M2 microscope (Zeiss) or Leica SP8 confocal microscopes. Image acquisition was done with 5x (for the tile scans), 10x, 20x, 40x and 63x objectives.

Images were processed and quantified with ZEN software (Zeiss), LAS (Leica), ImageJ software (NIH) or Cell Profiler (Broad Institute)^4^.

### Oxygen-induced retinopathy (OIR) model

A litter of P7 pups and their mother and/or foster mother were exposed to 75% oxygen in a semi-sealed oxygen chamber (ProOx 110 sensor and A-Chamber, Biospherix, Parish, NY) from P7 to P12, followed by room oxygen from P12 to P15-P17. Pups were sacrificed at indicated time point and the eyes were collected. To assess vascular permeability following OIR, P17 pups were warmed under a heat lamp and then 50 µl of green fluorescent microspheres (1% solution of 25 nm FITC conjugated microsphere Fluoro-MAX G25, Thermo scientific, Fremont, CA) were injected via tail vein using a 30-gauge insulin syringe while the mice were under temporary isoflurane anaesthesia. Following injection, microspheres were allowed to circulate for 15 minutes before the mice were once more placed under isoflurane anaesthesia. PBS was flushed through the vasculature via cardiac perfusion to remove excess microspheres, which had not extravasated followed by 4% PFA for tissue fixation. Eyes were enucleated and fixed in 4% PFA at RT for 30 minutes before the retinas were dissected and immunostained with isolectinB4 conjugated to Alexa647 and pERK1/2 as described.

To visualize Pald1 expression after OIR, eyes from P17 pups were collected and processed for lacZ staining as described below.

### Intravitreal injections and lacZ staining of retinas

10-week old female mice were anesthetized with isofluorane (AbbVie, Sweden). Prior to injection, the pupil was dilated with a drop of tropicamide (0.5% mydriacyl). A single injection of 1µg dog VEGF-A164 (1 µl injection volume) into the intravitreal space was done using a Hamilton syringe (Microliter #701RN with 34 gauge/25 mm/pst4 removable needle). The eyes were collected 72 h after injection and processed for lacZ staining.

For lacZ staining eyes were fixed in 0.4% PFA for 4 h at RT, retinas were dissected and permeabilized by washing three times for 20 min with detergent rinse (2 mM MgCl2, 0.01% sodium deoxycholate, 0.02% Nonidet P-40, PBS). Retinas were stained with 1 mg/ml x-gal (Promega) diluted in staining solution (detergent rinse containing 5 mM potassium ferricyanide, 5 mM potassium ferrocyanide) at 37°C overnight, protected from light. Retinas were washed twice for 10 min in detergent rinse, followed by two PBS washes and post-fixation with 4% PFA for 1 h at RT and mounted in Fluoromount-G (Southern Biotech). Images were acquired with AxioImager M2 (Zeiss).

### Quantitative real-time PCR

Eyes were collected from P4 and P5 pups and placed in RNA*later*^®^ solution (Ambion) immediately after collection. RNA was isolated using the RNeasy Micro Kit (Qiagen) and processed for quantitative real-time PCR as described previously^6^. The TaqMan Assays (Applied Biosystems) used: *Gapdh* (4352932E), *Ccnd1* (Mm00432359_m1, n=10/11).

