## Supplementary material for "Paladin is a PI(4,5)P_2_ phosphoinositide phosphatase that regulates endosomal signaling and angiogenesis": Suppl Figures 1-5

### Supplementary figure 1

**a**

```
Pald1_mus musculus (3)              311 LFSCQSGVGRT 323
PALD1_homo sapiens (3)              308 VFSCQMGVGRT 320

Pald1_mus musculus (4)              660 VFSCLSGQGRT 672
PALD1_homo sapiens (4)              657 VFSCLSGQGRT 669

PTPc, catalytic domain motif (M9)    43 VVHCSAGVGRT 55
VE-PTP (Ptprb)                     1901 VVHCSAGVGRT 1913
```

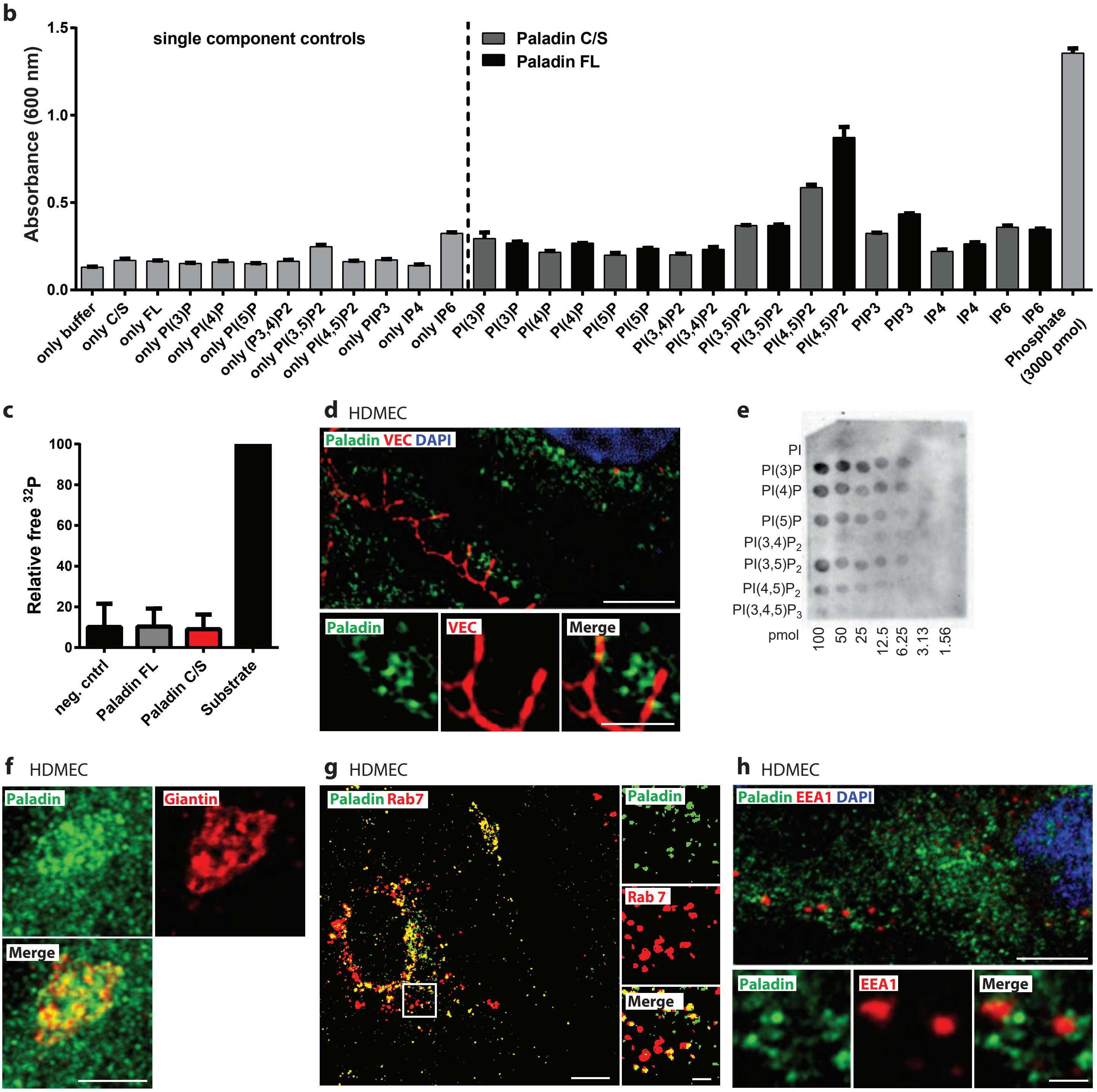

Supplementary Figure 2

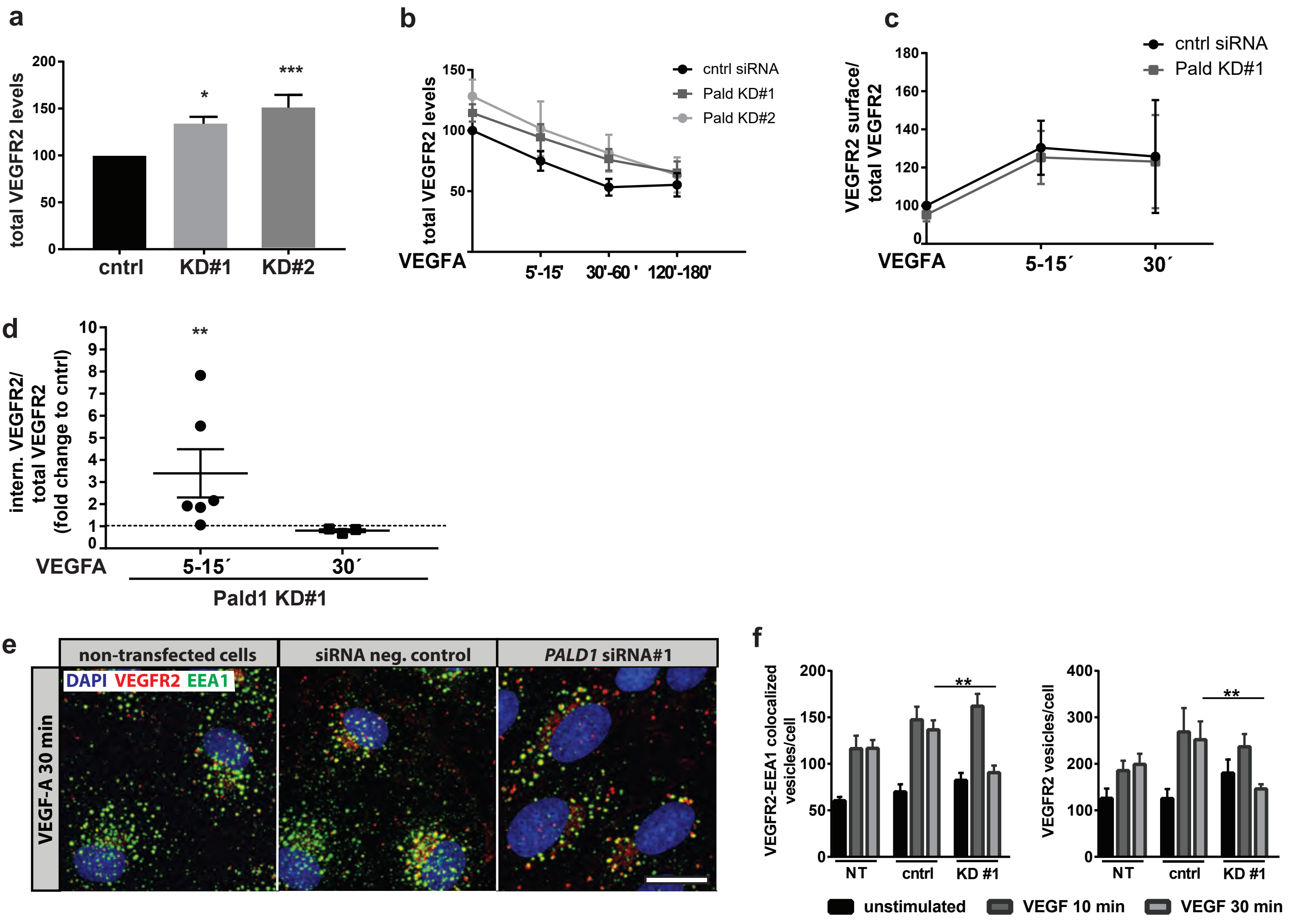

Supplementary figure 3

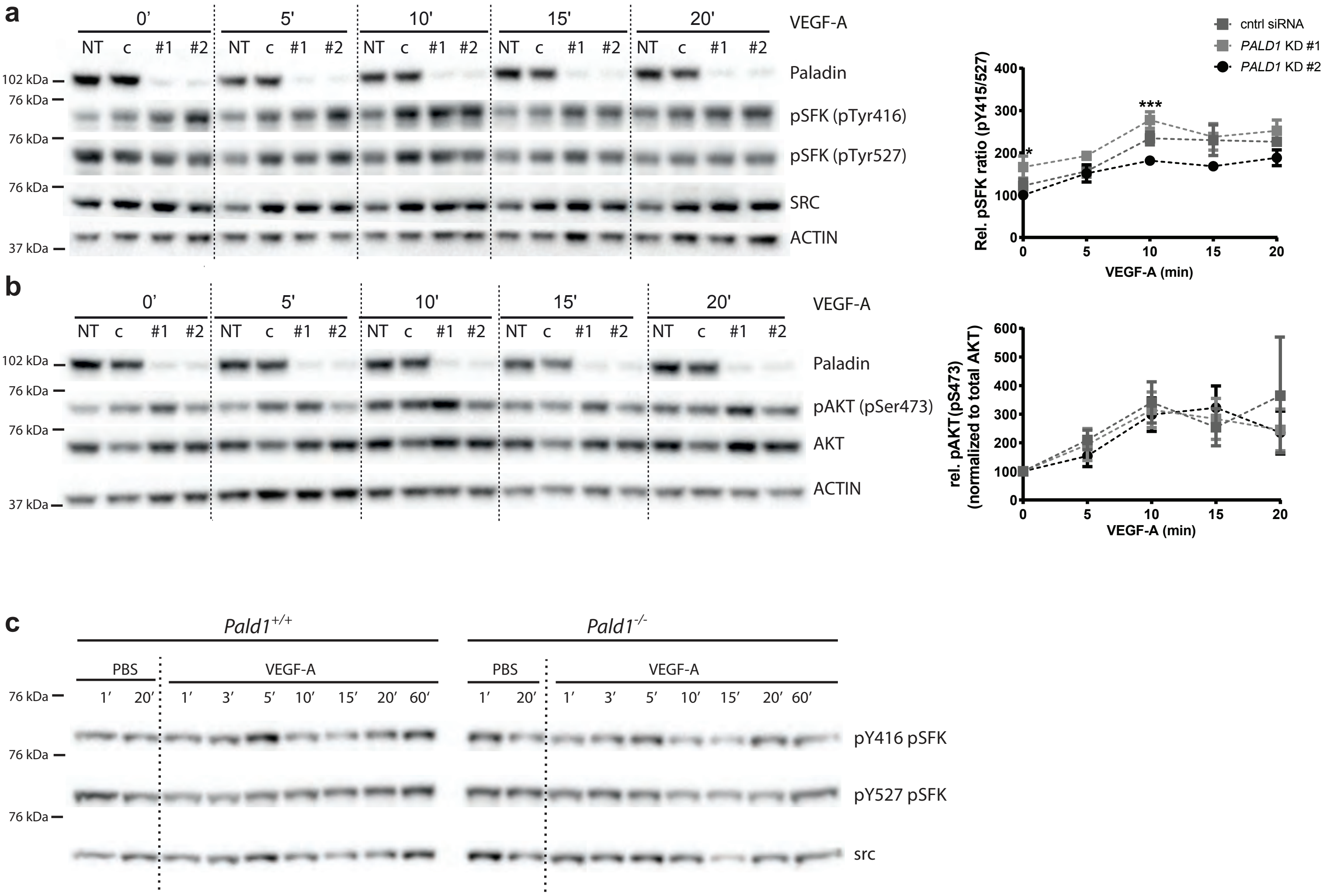

Supplementary figure 4

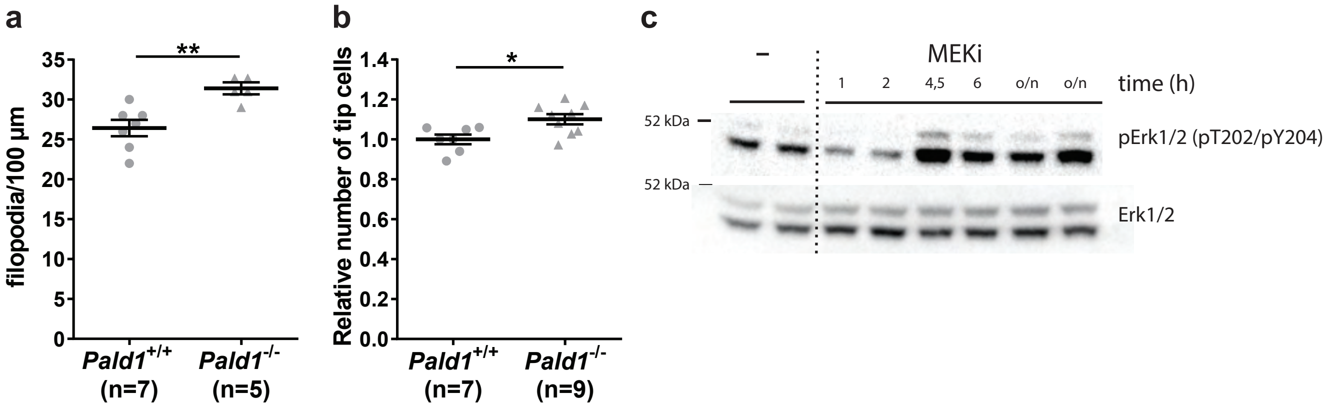

Supplementary Figure 5

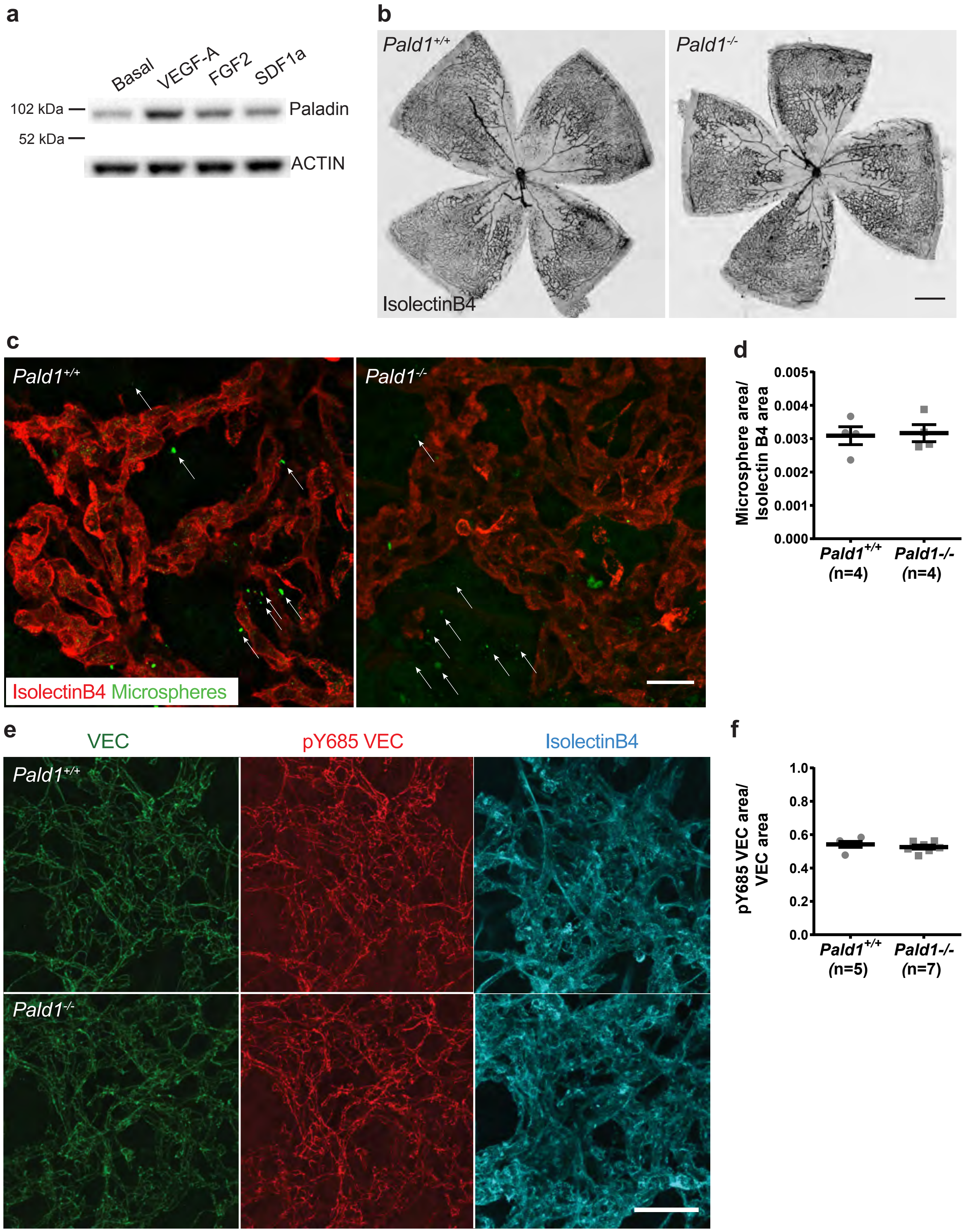
